# The immune response and microbiota profiles during co-infection with *P. vivax* and soil-transmitted helminths

**DOI:** 10.1101/2020.01.30.925032

**Authors:** Alice V. Easton, Mayra Raciny-Aleman, Victor Liu, Erica Ruan, Maria Fernanda Yasnot, Ana Rodriguez, P’ng Loke

## Abstract

Co-infection with soil-transmitted helminths (STH) and *Plasmodium* spp. parasites is a common occurrence in tropical low-income countries, but the consequences of this interaction remain poorly understood. Here, we performed a multi-omic analysis on peripheral blood and fecal samples from 130 individuals in Tierralta, Córdoba, Colombia who were infected with *P. vivax* alone (n = 33), co-infected with *P. vivax* and STH (n = 27), infected with STH alone (n = 39) or were infected with neither *P. vivax* nor STH (n = 31). In addition to Complete Blood Count (CBC) with differential, transcriptional profiling of peripheral blood samples was performed by RNA-Seq, fecal microbial communities were determined by 16S ribosomal RNA gene sequencing and circulating cytokine levels were measured by bead-based immunoassays. Differences in blood cell counts were driven primarily by *P. vivax* infection, including an increased percentage of neutrophils that was associated with a transcriptional signature of neutrophil activation in the blood. *P. vivax* infection was also associated with increased levels of IL-6, IL-8 and IL-10, and these cytokine levels were not affected by STH co-infection. Surprisingly, *P. vivax* infection was more strongly associated with changes in the microbiome than STH infection. Children infected with *P. vivax* exhibited elevated *Bacteroides* and reduced *Prevotella* and *Clostridiaceae*, but these differences were not observed in individuals co-infected with STH. We also observed that *P. vivax* parasitemia was higher in the STH-infected population. When we used machine learning to identify the most important predictors of *P. vivax* parasite burden from all measured variables, bacterial taxa were the strongest predictors of parasitemia levels. In contrast, circulating TGF-β was the strongest predictor of *T. trichiura* egg burden. This study provides unexpected evidence that the gut microbiota may have a stronger link with *P. vivax* than with STH infection.

## Introduction

Soil-transmitted helminth (STH) and malaria co-infection is a common occurance due to the geographical overlap of these infections (Mwangi et al. 2006). Both parasites can manipulate the host immune response to allow for their own persistence. Helminths can regulate the host immune system to prevent their elimination, which simultaneously protects the host against excessive inflammation (Maizels and Yazdanbakhsh 2003). Although this may be beneficial for the worm and the host in many cases, it also allows for other foreign antigens to remain hidden, thus possibly affecting the immune response to other pathogens (Hartgers and Yazdanbakhsh 2006). The effects of helminth co-infection are therefore complex, and epidemiological studies have often resulted in contradictory conclusions. One potential reason for contradictory results in the literature is that STH infection may render the host more susceptible to co-infection with other pathogens, but may also reduce the severity of morbidity resulting from inflammation in response to these other infections. For example, in response to malaria, the host produces high levels of pro-inflammatory cytokines that help control the parasite but also contribute to pathology (Moxon et al. 2019). In endemic areas, repeated infections lead to the development of partially protective immunity to malaria, which frequently results in persistent infections with low levels of parasites (Bousema et al. 2014).

Several studies have found an association between malaria severity and the presence of STH. However, the direction of the association is dependent on the study and the species of STH in the coinfection. One study from Senegal showed that the children who tested positive for *Ascaris, Ancylostoma* or *Trichuris* were more susceptible to *P. falciparum* infection than their uninfected peers (Spiegel et al. 2003). However, another study also in Senegal showed that *Schistosoma haematobium* infection was associated with lower *P. falciparum* densities (Briand et al. 2005). When it comes to pathogenesis, a study in Thailand found that *Ascaris* infection was protective against cerebral malaria (as opposed to uncomplicated malaria cases), suggesting that that *Ascaris* in particular might protect against morbidity resulting from malaria infection (Nacher et al. 2000). It is hypothesized that people infected with helminths are more susceptible to *P. falciparum* infection, but experience severe morbidity from these infections less frequently (Druilhe et al. 2005; Degarege et al. 2016). In mouse models of coinfection, there is a similar confusion over the contrasting effects of helminths, depending on the nature of the rodent malaria model (Knowles 2011). Variation in the gut microbiota between individual mammalian hosts may further confound some of these complex interactions.

There is now evidence that the composition of the gut microbiota in the mammalian host can affect disease severity and protect against infection with malaria parasites (Ippolito et al. 2018). In mice, *Plasmodium* infection can affect the composition of the gut microbiota, changing their susceptibility to other infections or to malaria-induced pathology (Taniguchi et al. 2015; Mooney et al. 2015). Different gut bacterial communities of mice from different vendors were responsible for differences in parasite burden and mortality after infection with *Plasmodium* (Villarino et al. 2016). These studies in mouse models indicate that malaria can induce changes in the gut microbiome of the host, but also that the composition of the host microbiome can modulate the development of pathology induced by malaria. In human subjects, minimal differences in the composition of the gut microbiome were found in children before and after malaria followed by artesunate treatment (Mandal et al. 2019), indicating important differences with mouse models of infection in this respect. However, similarly to mice infections, a human-subjects study from Mali suggested that the gut microbiome may affect susceptibility to *P. falciparum* infection; individuals who were at a lower risk of being infected with *P. falciparum* had a significantly higher proportion of *Bifidobacterium* and *Streptococcus* in their gut microbiome (Yooseph et al. 2015).

The effects of helminths on the gut microbiota have also been increasingly documented (Cortés et al. 2019). By characterizing the microbiota of a group of indigenous Malaysians known as the Orang Asli using 16S rRNA gene sequencing, we previously found that STH-infected individuals have greater microbial diversity than uninfected individuals (Lee et al. 2014). Deworming treatment reduces microbial diversity and shifts the community balance by increasing *Bacteroides* abundance and reducing *Clostridiacae* in this population (Ramanan et al. 2016). Furthermore, the parasite burden of *Trichuris* infection has one of the largest effect sizes on microbial variation (Lee et al. 2019). Nonetheless, every study population has unique characteristics, and in other study sites, such trends have not been observed (Martin et al. 2018; Cooper et al. 2013). Large-intestinal-dwelling helminths, such as *Trichuris trichiura*, may have different effects on the microbiota than small-intestinal and tissue-dwelling helminths (Cortés et al. 2019). Additionally, the effects of the anti-helminthic drugs on the microbiota is still not definitively understood. Finally, how helminths, malaria and the microbiota interact to influence the immune response remains unclear.

Here, we examine the association of STH and *P. vivax* co-infection with the gut microbiota and immune response through a cross-sectional study based in Colombia. We identified the most important predictors of both *T. trichiura* and *P. vivax* parasiemtia levels using machine learning models to integrate the various multi-omic measurements. Surprisingly, the gut microbiota may have a stronger relationship with *P. vivax* infection than STH infection, since multiple bacterial taxa were identified to be the strongest predictors of parasitemia levels in *P. vivax* infected individuals.

## Results

To study the effects of *P. vivax* and STH co-infection, we collected blood and stool samples from 130 children aged four to sixteen, living in Tierralta, Córdoba, Colombia. Healthy children infected (n = 39) or not (n = 31) with STH were confirmed to be PCR negative for *P. vivax* in the blood. Children with acute *P. vivax* malaria (n = 60) were split into those who were infected with any STH (n = 27), and those who were not (n = 33). These two groups had comparable proportions of female and indigenous individuals, and similar age distributions (Table 1). Notably, children co-infected with STH had higher *P. vivax* parasitemia measurements (Table 1, Figure 1, p=0.04 by Wilcoxon signed-rank test). This may indicate that STH infection makes the host more permissive to *P. vivax* parasites.

**Figure 1.**
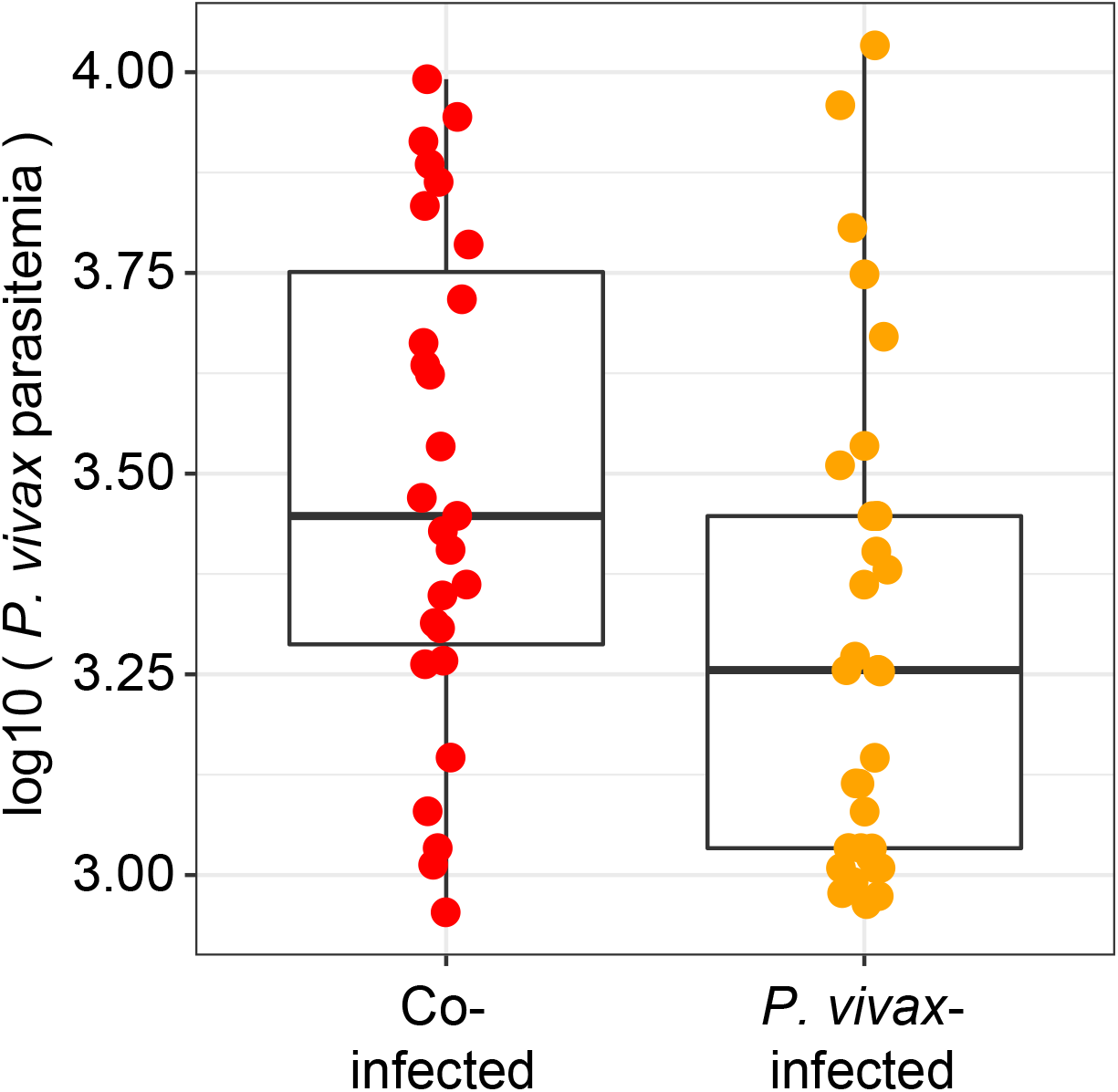
*P. vivax* parasitemia is higher in STH co-infected individuals. Boxplot showing *P. vivax* parasitemia levels of children with both *P. vivax* and STH infection (red dots), and children with *P. vivax* infection only (orange dots). *P. vivax* parasitemia is significantly higher in the individuals who are co-infected with STH (p=0.04 by Wilcoxon signed-rank test).

**Table 1.** Characteristics of the study population. The study population included four groups of children: those infected with *P. vivax* alone, infected with STH alone, infected with both *P. vivax* and STH, or uninfected with any of *P. vivax* or STH. Children infected with *P. vivax* (with or without STH infection) were enrolled in the hospital before receiving treatment. Children uninfected with *P. vivax* (with or without STH infection) were enrolled from a school within the hospital’s catchment area.

| Group | Uninfected | *P. vivax* -<br>infected | STH -<br>infected | Co-infected | Differences between groups |
| --- | --- | --- | --- | --- | --- |
| Participants | 31 | 33 | 39 | 27 |  |
| Female | 16 | 18 | 13 | 8 | No sig. diffs. by two-proportion z-test |
| Indigenous | 0 | 11 | 0 | 7 | Hospital-collected populations similar<br>(two-proportion z-test) |
| Age | 9.1±2.9 | 12.8±3.2 | 10.6±2.2 | 12.0±2.98 | *Differences by ANOVA. Sig. diffs*<br>*between all pairs except P. vivax and*<br>*Co-infected* |
| *P. vivax* parasitemia | 0 | 2554±2346 | 0 | 3901±2660 | *Differences by ANOVA. p=0.04*<br>*between P. vivax and Co-infected* |
| Number infected<br>with *T. trichiura* | 0 | 0 | 36 | 25 | No sig. diffs. by two-proportion z-test |
| *T. trichiura* egg<br>count | N/A | N/A | 38±44 | 50±88 | No sig. diffs. by Mann-Whitney test |
| Number infected<br>with *A. lumbricoides* | 0 | 0 | 11 | 4 | No sig. diffs. by two-proportion z-test |
| *A. lumbricoides* egg<br>count | N/A | N/A | 2115±3825 | 347±659 | No sig. diffs. by Mann-Whitney test |
| Number infected<br>with Hookworm | 0 | 0 | 3 | 11 | *Sig. diff. by two-proportion z-test*<br>*(p=0.001)* |
| Hookworm egg count | N/A | N/A | 295±502 | 37±65 | No sig. diffs. by Mann-Whitney test |
| Location collected | School | Hospital | School | Hospital |  |

The frequencies of infection with *Trichuris trichiura* and *Ascaris lumbricoides* were similar between children who were co-infected with *P. vivax* versus those who were not. Although more individuals had hookworm infections in the *P. vivax* co-infected group (n = 11) than the *P. vivax* negative group (n = 3), hookworm prevalence was relatively low (21%), compared to *T. trichiura* (92%). There were no significant differences between the egg counts for each of these three STH between children infected with STH versus those co-infected with STH and *P. vivax* (Table 1).

A Complete Blood Count with differential (CBC w/ diff) analysis of the blood for the 130 children in this study provided data on cellular composition and other blood parameters. A PCA plot based on all the measurements from the CBC w/ diff shows that children infected with *P. vivax* alone or with STH co-infection largely cluster separately from those uninfected with *P. vivax* (Figure 2A). Hence, infection with *P. vivax*, but not with STH, significantly alters multiple blood parameters. Individuals with *P. vivax* infection diverge from individuals without *P. vivax* infection along the PC1 axis (Figure 2A). This indicates that PC1 can explain an important component of the variance that separates *P. vivax* infected and uninfected individuals. Identification of factors with negative loading on PC1 (Figure 2B) indicates that there is a negative association between *P. vivax* infection and platelets/hematocrit (Figure 2B and 2C), which is consistent with the known consequences of *P. vivax* infection (Coelho et al. 2013; Leal-Santos et al. 2013). Additionally, identification of factors with a positive loading on PC1 (Figure 2B) indicates that there is a positive association between *P. vivax* infection and the percentage of neutrophils/monocytes in the blood (Figure 2B). Indeed, platelets and hematocrit are significantly lower in individuals infected with *P. vivax*, regardless of STH infection (Figure 2C) and the percentage of monocytes and neutrophils are elevated in individuals with *P. vivax* infection (Figure 2C). Boxplots for the remaining six variables can be found in Figure S1.

**Figure 2.**
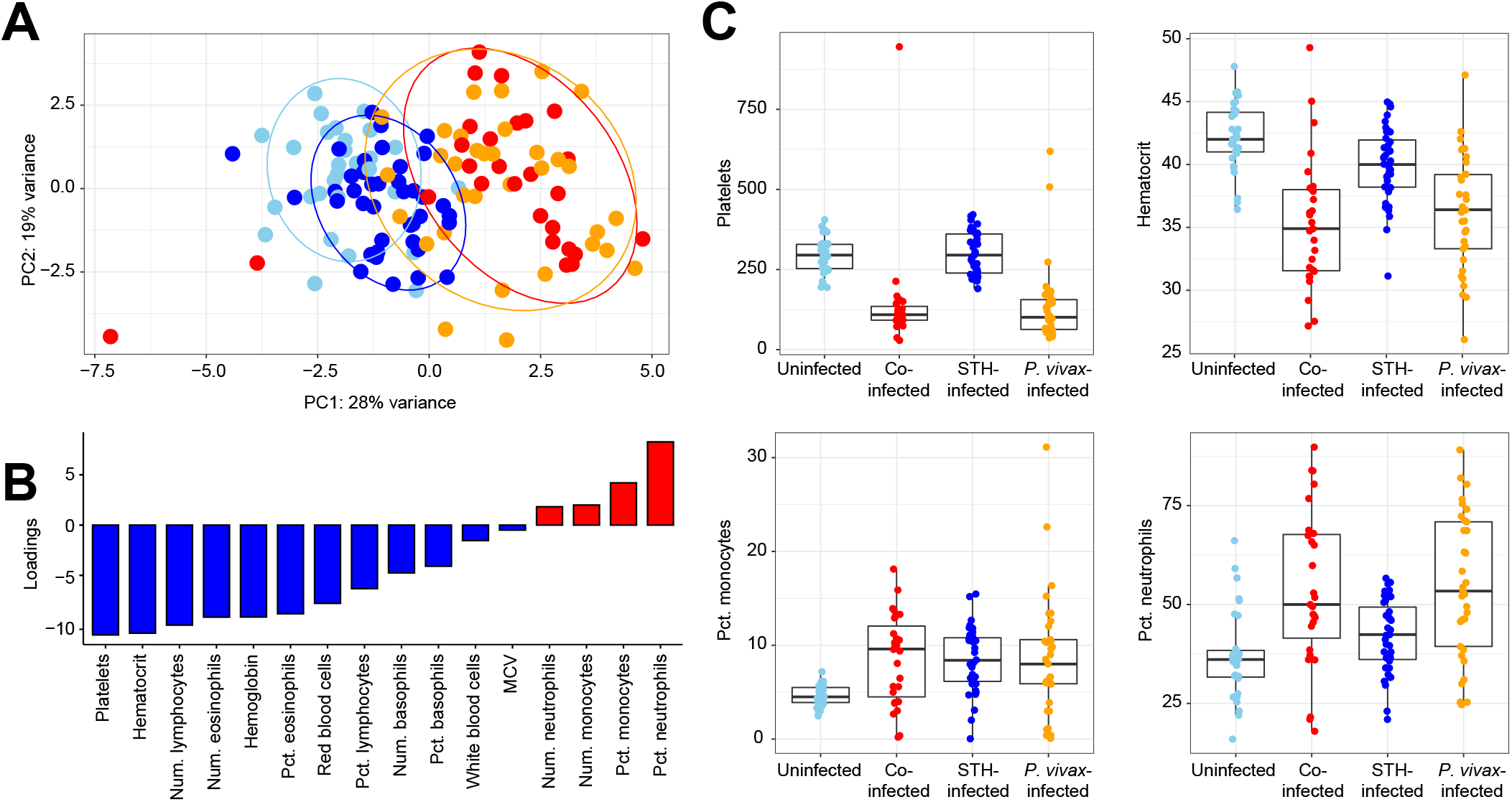
Complete blood count with differential (CBC w/ diff) analysis can distinguish individuals with *P. vivax* infection but not with STH infection. (A) Principal components analysis (PCA) based on results from CBC w/ diff analysis. Each point represents one child and is color-coded by infection status, as throughout the manuscript. Children co-infected with *P. vivax* and STH are shown in red, children with *P. vivax* alone are shown in orange, children with STH alone are shown in dark blue, and children uninfected with either parasite are shown in light blue. Ellipses show the area covering 90% of the samples from each group. (B) The factors loading for PC1 reflect the amount of variance shared by these parameters (either negatively in blue, or positively in red) with the PC1 values. (C) Boxplots of the two variables most negatively associated with PC1 (platelets and hematocrit) as well as the two variables most positively associated with PC1 (the percentage of monocytes and neutrophils) are shown with the same color scheme. Boxplots for the remaining six clinical variables can be found in Figure S1.

To examine the relationship between specific cytokines and infection status, we measured circulating levels of 13 cytokines in plasma samples. A PCA plot based on these variables shows that most individuals had similar levels of these cytokines, except for some individuals with *P. vivax* infection either with (red dots) or without (orange dots) STH co-infection (Figure 3A). IL-6, IL-8, IL-10 and IL-4 are variables that contributed the most to the positive loading along the PC1 axis (Figure 3B). These four cytokines were elevated in individuals infected with *P. vivax*, with or without STH coinfection (Figure 3C).

**Figure 3:**
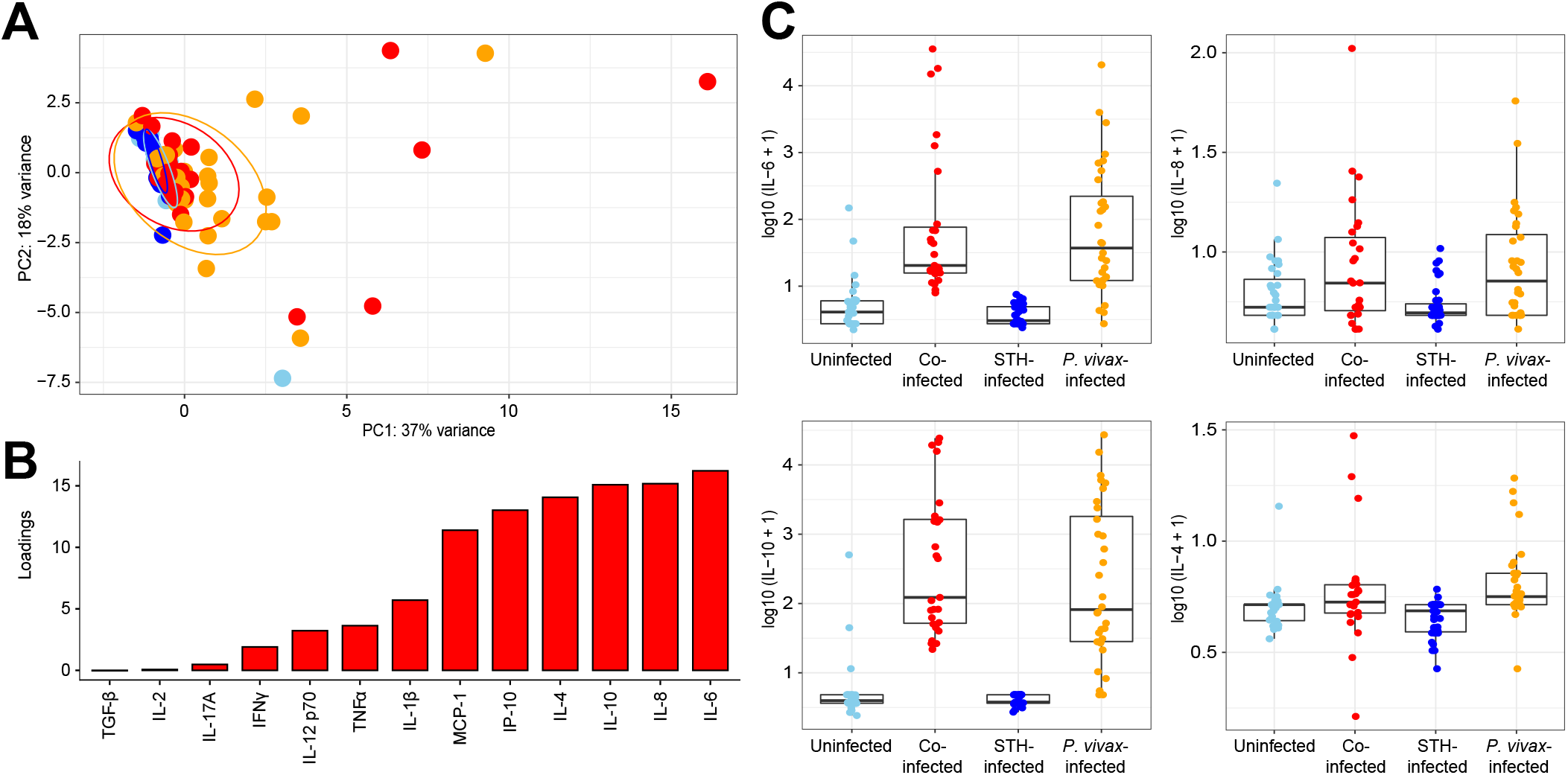
Elevation of plasma cytokine levels in some individuals infected with *P. vivax.* (A) Principal components analysis (PCA) of 13 different circulating cytokine levels measured by bead-based immunoassays. Each point represents one child and is color-coded by infection status, including children co-infected with *P. vivax* and STH (red), children with *P. vivax* alone (orange), children with STH alone (dark blue), and children uninfected with either parasite (light blue). Ellipses show the area covering 90% of the samples from each group. (B) The factors positively loading for PC1 reflect the amount of variance shared by these parameters with the PC1 values. (C) Boxplots of the four variables (IL-6, IL-8, IL-10 and IL-4) most positively associated with PC1.

To identify associations between the gut microbial communities and infection status, we performed 16s rRNA gene sequencing on stool samples collected from all children in the study. A Principal Coordinates (PCoA) plot was constructed to map each sample onto two-dimensional space, using Jaccard distances (Figure 4A). Most individual samples cluster together, though some samples from children with *P. vivax* infections alone (without STH co-infection, shown in orange) cluster separately along the first axis. When comparing the microbiota composition of children with *P. vivax* infection alone (orange) or with STH co-infection (red), *Bacteroides* was elevated in individuals without STH co-infection. *Prevotella copri, Prevotella* and *Clostridiaceae* were elevated in co-infected children (Figure 4B and Figure 4C). *Prevotella copri*, Prevotella and Clostridiaceae were less abundant in the samples from the *P. vivax*-infected group, and *Bacteroides* was more abundant in samples from the *P. vivax*-infected group than in any of the other groups. These results indicate that differences in gut microbial composition are associated with *P. vivax* infection in some individuals, but none of the *P. vivax* infected individuals who were co-infected with STH show similar alterations in microbial composition.

**Figure 4:**
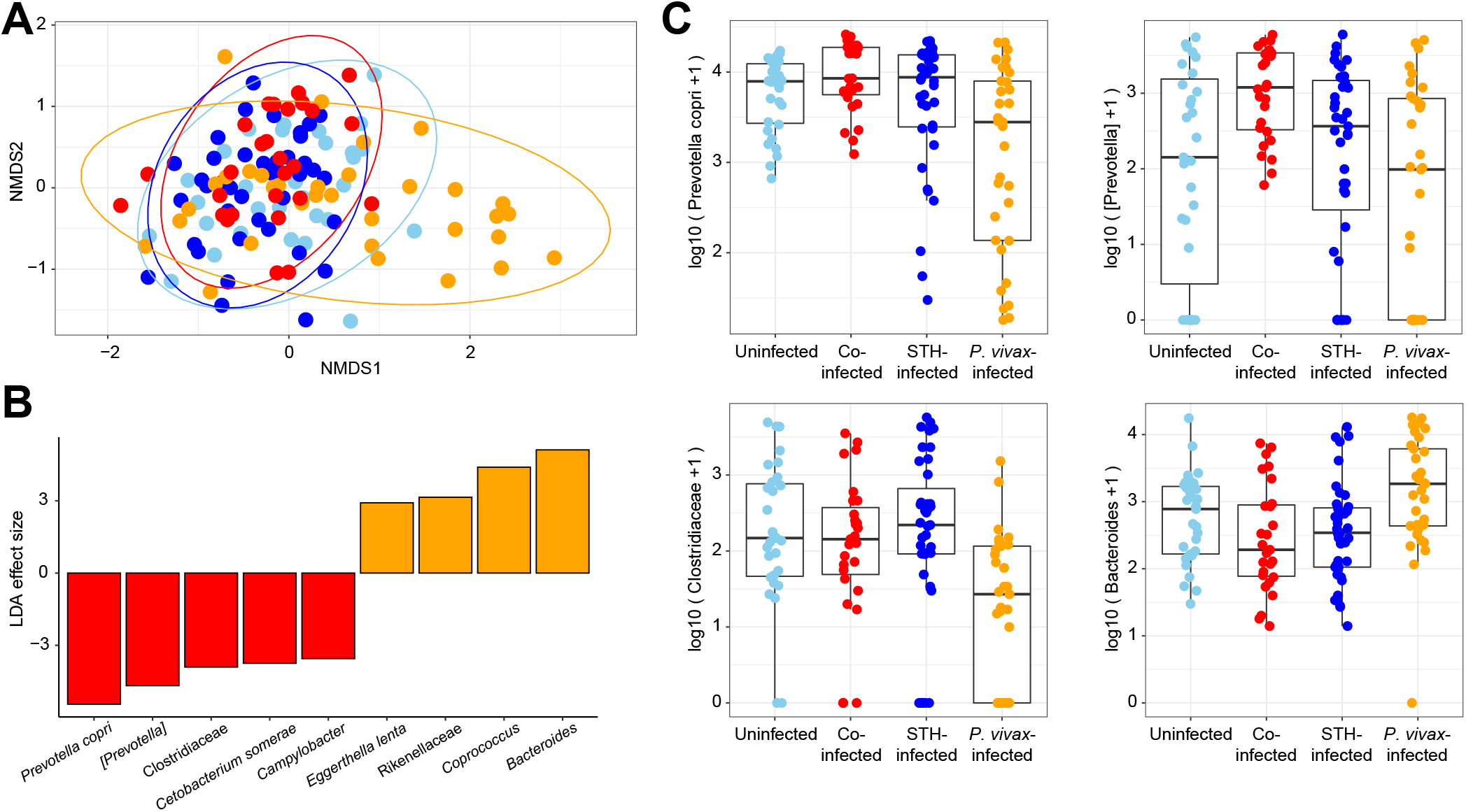
Effects of STH co-infection on the microbiota of *P.* vivax-infected patients. (A) An NMDS plot based on Jaccard distance between stool microbiota samples (based on 16s rRNA gene sequencing), colored according to STH/P. *vivax* infection status. Some individuals with *P. vivax* infections (orange) (without STH co-infection(red)) have a distinct microbiota, and cluster separately along PC1. (B) LDA effect sizes calculated using LEFSe (Segata et al. 2011) are shown, comparing samples from individuals with *P. vivax* alone versus those co-infected with *P. vivax* and STH. Microbes that are significantly (p<0.01) elevated in individuals with *P. vivax* alone are shown in orange, and microbes that are significantly elevated in individuals with co-infection with both *P. vivax* and STH are shown in red. (C) Boxplots for the three top microbes enriched in samples from co-infected individuals, and the top microbe enriched in samples from *P. vivax*-infected individuals.

We next performed transcriptional analysis by RNA-Seq of the total blood samples that were collected into PAXgene tubes from enrolled children. A PCA plot based on the total RNA-Seq dataset shows that most samples have similar transcriptional profiles. There are some individual outliers, but these samples come from all four groups of children (Figure 5A). DESeq analysis identified 30 genes that were significantly upregulated in children with *P. vivax* infection (Figure 5B). Functional enrichment analysis of these 30 genes using Panther (Mi et al. 2019) indicated that many of these genes are involved in neutrophil function (Figure 5C). Hence, *P. vivax* infection appears to be associated with an increase in neutrophil activation in the peripheral blood. As noted above, CBC w/diff analysis had indicated that the percentage of neutrophils was elevated in individuals with *P. vivax* infection (Figure 2C). However, the absolute number of neutrophils was similar across all groups (Figure S1B). Hence, this neutrophil signature may reflect an increased activation state of blood neutrophils and not just an increase in cell number. DESeq analysis comparing individuals with STH infection versus uninfected individuals (and comparing individuals co-infected with *P. vivax* and STH versus infected with *P. vivax* alone) did not identify any genes that were significantly different, indicating that the transcriptional effects of helminth infection on the blood cells are relatively minor.

**Figure 5:**
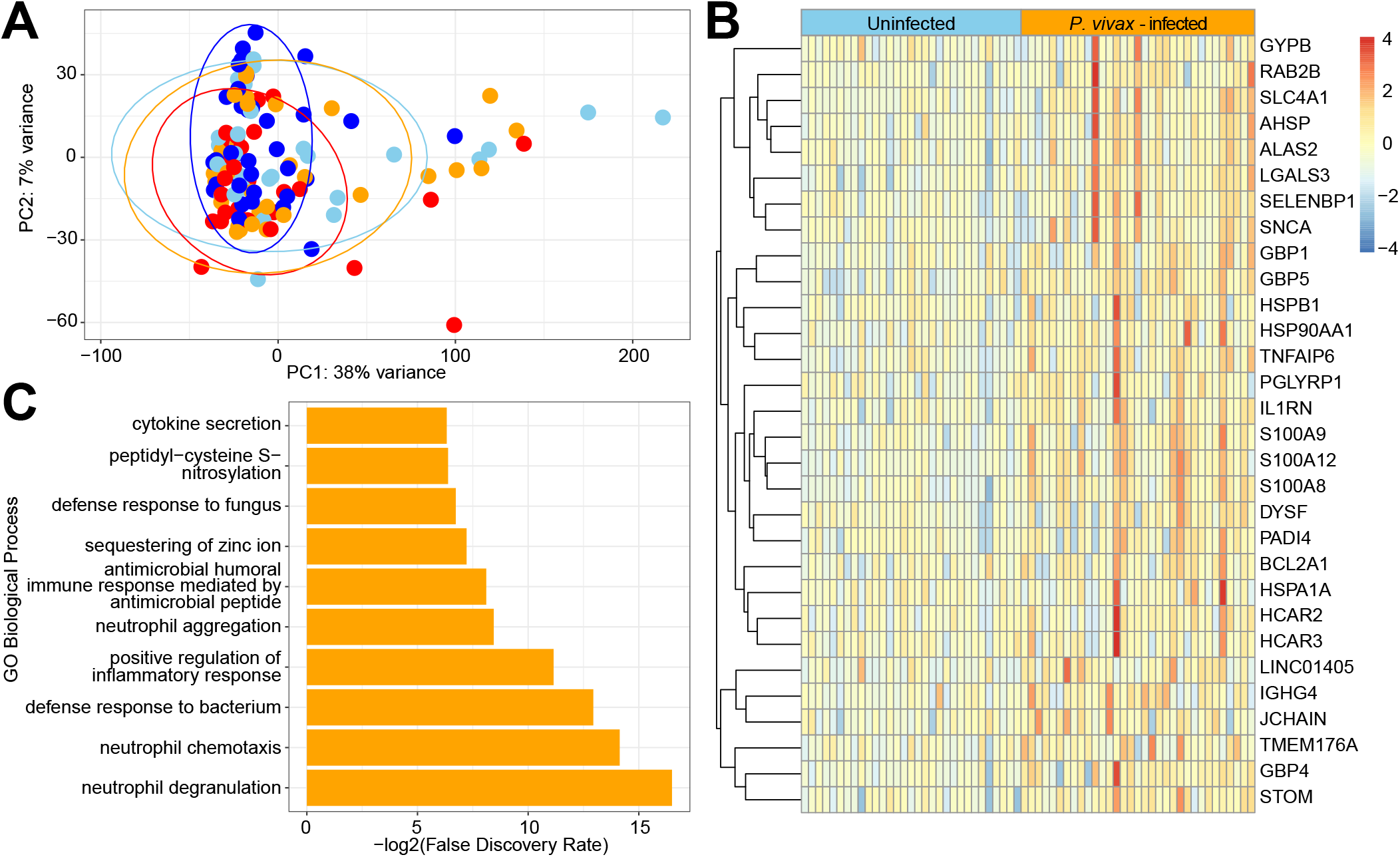
RNA-Seq of the peripheral blood identifies genes upregulated by *P. vivax* infection, but not by STH infection. (A) PCA plot based on the top 50% most variable genes with at least 400 reads across all samples. Ellipses show the area covering 90% of the samples from each group, showing the almost complete overlap between groups. (B) Differential abundance analysis by DESeq identifies 30 genes that are upregulated in *P. vivax*-infected individuals, as compared to uninfected individuals (with an adjusted p-value <0.05 and a log2 fold change >=1). (C) Biological processes over-represented in these 30 genes relative to all genes in the *Homo sapiens* GO Ontology database (10/8/19), using Fisher’s Exact test in PANTHER (Mi et al. 2019). The top ten specific subclasses (from hierarchically sorted output) are shown, based on lowest FDR.

Having assembled a heterogenous set of blood and stool measurements from this study, we wanted to integrate the data and determine which variables were the most important predictors of the four separate groups by feeding all the variables into a random forest model. Since we were most interested in the effects of co-infection, we focused on the differences between individuals with *P. vivax* infection, versus those with both *P. vivax* and STH co-infection. When the variables were sorted by importance to the predictive model (the mean decrease in Gini when the variable was omitted from the model), 8/10 of the most important variables were microbes (Figure 6A). As expected, some of these microbes overlapped with those identified by LDA (Figure 4): *[Prevotella], Prevotella, Bacteroides*, Clostridiaceae and *Cetobacterium.* Other important variables include one gene (NDUFA6, NADH:Ubiquinone Oxidoreductase subunit A6, not identified as differentially expressed by DESeq2) and *P. vivax* parasitemia. As noted above, STH co-infection was associated with higher *P. vivax* parasitemia (Table 1 and Figure 1). However, it should be noted that the out-of-box error rate for this model was 38% and a random model would have an error rate of approximately 50%. Hence, the large number of variables we measured only provided a small amount of information that could differentiate between STH co-infected and *P. vivax* infected individuals. Nevertheless, the top predictors identified by the model both indicate that the gut microbiota may be an important contributor to further understanding the effects of *P. vivax* and STH co-infection, and identified microbes found (by LDA analysis and individual inspection in boxplots) to be different in the *P. vivax* infected group when compared to all other groups.

**Figure 6:**
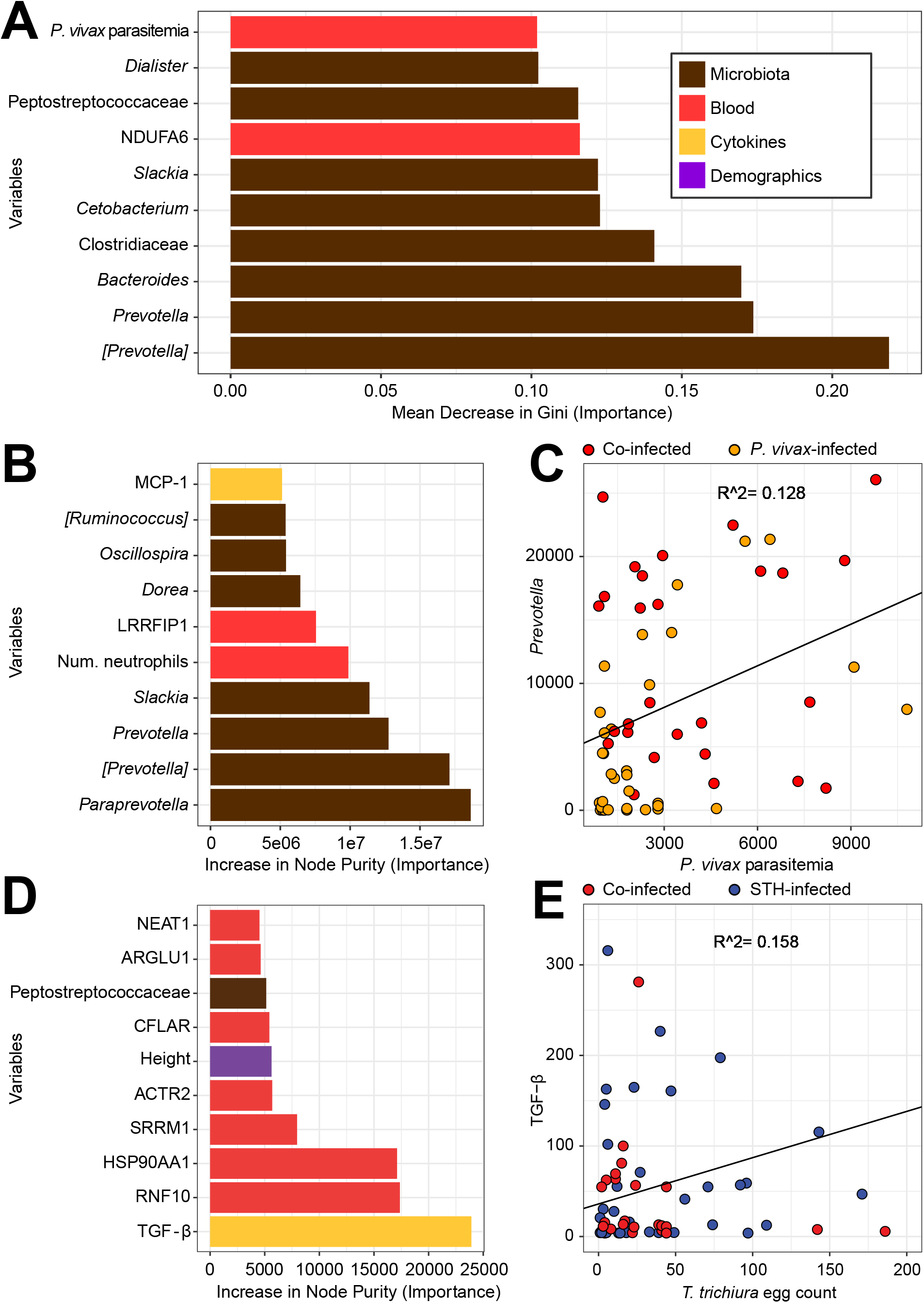
Integrative analysis of heterogeneous datasets using random forest models to identify the strongest predictors of co-infection and parasite burden. (A) Random forest model selected eight microbes (brown) in top ten predictors of whether a *P.* vivax-infected child is also infected with STH. Bars represent the mean decrease in Gini when a variable is removed from the model; a larger decrease means that the variable is more different between the individuals with *P. vivax* alone versus both *P. vivax* and STH infection. Bars are color-coded based on the type of variable: microbes measured by 16s rRNA gene sequencing are in brown, genes measured by RNA-Seq are in red, cytokines are in yellow and demographic variables from a questionnaire are in purple. The model included 4046 variables: 38 microbial genera, 3907 genes, 85 measurements from blood tests including CBC with diff and cytokine levels and 16 variables from the demographic questionnaire. (B) Random forest model selects seven microbes in top ten predictors of *P. vivax* parasitemia. In this continuous-outcome model, the increase in node purity represents the importance of the variable to the model; higher numbers mean the variable is more important for predicting *P. vivax* parasitemia. C) To examine one of these important variables, we created a scatterplot, which shows that *Prevotella* is correlated with *P. vivax* parasitemia (r^2^ = 0.13, p-value = 0.005, among those infected with *P. vivax).* Samples from children infected with *P. vivax* are shown in orange, and samples from children co-infected with STH are shown in red. D) Random forest model selects TGF-β as top predictor of *T. trichiura* egg count. Other variables that are predictive of egg burden include several genes (from RNA-Seq results), the child’s height, and one microbe. E) TGF-β is correlated with *T. trichiura* egg count (r^2^ = 0.16, p-value = 0.002, among those infected with *T. trichiura).* Samples from children infected with STH are shown in blue, and samples from children coinfected with *P. vivax* are shown in red.

Next, we wanted to determine what variables are important predictors of parasite burden for *P. vivax* and *Trichuris trichiura.* At 92% prevalence, *T. trichiura* was by far the most common STH found in this community. First, a random forest model was run with the continuous outcome of *P. vivax* parasitemia (Figure 6B). Strikingly, some of the same microbes identified in the categorial classification above (Figure 6A) were also identified as important predictors of *P. vivax* parasitemia levels. Of the 10 most important predictors of parasitemia, seven were bacterial taxa, highlighting the potential importance of gut bacterial communities in *P. vivax* infection. Microbes that were predictive both of whether a person was coinfected with STH, and of *P. vivax* parasitemia, included *[Prevotella], Prevotella* and *Slackia.* The number of neutrophils was also predictive of *P. vivax* parasitemia, which is consistent with the other results (Figure 2 and Figure 5) indicating a role for neutrophils in individuals infected with *P. vivax*. When we plotted *Prevotella* read counts against *P. vivax* parasitemia (Figure 6C), the coefficient of correlation was low (r2=0.128), but the p-value for this correlation was significant at 0.005.

When we ran the random forest model to predict *T. trichiura* egg count, the strongest predictor was TGF-β (Figure 7D). When we plotted TGF-β levels against *T. trichiura* egg count (Figure 7E), the coefficient of correlation was low (r2=0.158), but the p-value for this correlation was significant at 0.002 (Figure 7E). Notably, although the parameters measured by CBC w/diff (Figure 2) and plasma cytokine measurements (Figure 3) appeared to be better at distinguishing individuals from separate groups, these parameters were mostly poor predictors of parasite burdens and co-infection status (with the exception of TGF-β and neutrophil numbers). Of the 10 most important predictors for *T. trichiura* egg count, seven parameters were transcript levels from the RNA-Seq analysis of whole blood. Hence, we found (to our surprise) that stool sample analysis for bacterial communities identified more predictors of *P. vivax* parasitemia, whereas blood sample analysis identified more predictors of *T. trichiura* egg count, which is the opposite of where the parasites reside.

## Discussion

In this study, we took a multi-omic approach towards investigating the effects of soil-transmitted helminths and *P. vivax* co-infection. To our surprise, we found that the gut microbiota communities had a stronger association with malaria infection, than with STH infections. We had previously reported, in both cross-sectional (Lee et al. 2014) and longitudinal studies on the Orang Asli in Malaysia (Ramanan et al. 2016), that *T. trichiura* infection impacted microbial diversity and the composition of the microbiota in infected individuals. In contrast, this current study shows a minimal effect of *T. trichiura* infection compared to uninfected individuals. This suggests that the environment in which the microbiota and STH exist (all the other co-factors, such as the diet, age, lifestyle, and other previous and current infections of the human host) may determine whether or not STH affect the gut microbiota. This could explain why the effects of STH on the gut microbiome have not been consistent between studies, even when examining the same STH species (Easton et al. 2019; Rosa et al. 2018; Martin et al. 2018; Cooper et al. 2013). Thereby we conclude that the effects of STH infections on the microbiota are dependent on the context of infection.

Here we observed an association of acute *P. vivax* infections with microbiome composition; however, the direction of the causal relationship between malaria and the microbiome is still not clear. It could be interpreted as changes in the intestinal microbiota communities being induced by infection with *P. vivax*, or as the result of individuals with particular microbiome characteristics being more susceptible to *P. vivax* infections. Previous longitudinal studies observed that there are minimal effects of *P. falciparum* infection on the microbiome composition (Mandal et al. 2019), but that the intestinal microbiota composition is associated with the prospective risk of *P. falciparum* infection (Yooseph et al. 2015). Taken together, these studies suggest that the pre-existing microbiota composition may affect the susceptibility to malaria.

The effect of STH infections in malaria development and pathogenesis is a highly relevant clinical issue because of the major overlap between these two parasites worldwide (Brooker et al. 2007). We found here that STH-infected individuals have higher *P. vivax* parasitemia (Figure 1). Previous studies in Southeast Asia (Burdam et al. 2016) and Africa (Tuasha et al. 2019; Degarege et al. 2012; Babamale et al. 2018; Efunshile et al. 2015) also reported that helminth co-infection had a positive association with occurrence of malaria caused by *P. falciparum* or *P. vivax* infections. While STH may render co-infected people more susceptible, it does not appear to cause increased severity of symptoms (Degarege et al. 2009; Hürlimann et al. 2019). In fact, there are also reports that helminth co-infection may protect against certain aspects of pathogenesis, such as *P.* vivax-induced anemia (Melo et al. 2010). These effects are probably mediated by immune modulation induced by helminth infections, which may alter the subsequent response to malaria, thus resulting in higher tolerance to disease (Frosch and John 2012).

We also observed an increase in the percentage of neutrophils among white blood cells and in the expression of neutrophil-specific genes in *P. vivax* infections, regardless of STH co-infection, confirming previous evidence that neutrophil activation is an important response to malaria (Aitken et al. 2018). Indeed, a strong neutrophil response with the formation of extracellular traps is observed in acute *P. falciparum* patients (Knackstedt et al. 2019). However, a previous study (Vallejo et al. 2018) described a decrease in neutrophils and neutrophil specific genes in *P. vivax* infections, but this was only observed in first-time infections, possibly accounting for the differences with our results.

In this study, we found that of all the multi-omic measurements that we performed, the level of circulating TGF-β was the variable most predictive of *T. trichiura* egg burden. Just based on this association, it is not clear whether TGF-β is important for worm expulsion or colonization. However, the relationship between TGF-β and *Trichuris* infection has been established in the murine model of *Trichuris muris* infection (Worthington et al. 2013). In this model, TGF-β was shown to act on CD4+ T cells to promote chronic infection instead of worm expulsion. Antibody-mediated blockade of TGF-β activity could protect mice from infection, indicating that the increased TGF-β levels may be beneficial for the parasite to colonize its host. One possibility may be through the induction of Foxp3+ CD4+ Tregs that promote chronic infection by suppressing protective immunity. A previous study on African children found that both *Trichuris trichiura* and *Ascaris lumbricoides* infection burden was associated with IL-10 and TGF-β production from peripheral blood cells (Turner et al. 2008). Notably, this study also showed that there was a negative relationship between IL-10 and TGF-β production and the ability of these cells to produce TH1 or TH2 cytokines against worm and bacterial antigens. In a different study, a healthy volunteer who ingested eggs from the porcine parasite *Trichuris suis* also expressed higher levels of TGF-β in the ascending and transverse colon after infection, indicating a direct link between infection and expression of TGF-β in humans (Williams et al. 2017). Finally, there is also evidence from human genetic association studies that TGF-B1 could be important in *T. trichiura* infection. Polymorphisms in the *TGF-B1* gene that affect allergy and asthma are also associated with increased susceptibility to *Trichuris trichiura* infection, as well as increased IL-10 production (Costa et al. 2017). Overall, our study provides additional evidence that the link between TGF-β and *Trichuris* infection may be one of the most important immunological relationships between the host and this parasite.

An important caveat of our analysis is that for the random forest regression model we used, the percent of variance explained was only −10.32 (for *P. vivax* parasitemia) and −23.35 (for *T. trichiura* egg burden), suggesting severely limited power to predict parasitemia from other types of data. For predicting co-infection membership, the model correctly classified 26/33 samples as *P.* vivax-infected, but incorrectly classified 16/27 co-infected samples as *P. vivax*-infected. This indicated that the large set of multi-omic parameters we measured only provided a small amount of information that can be used to differentiate between samples from these two groups. These models were more useful for identifying specific parameters that were interesting in the datasets and for generating hypotheses, than for any substantive predictive value. Much larger sample sizes would have to be collected to improve upon the current studies.

We believe that the current study provides additional evidence that the gut microbiota may play an important role in *Plasmodium* infections and contribute to the complexities of co-infection with STH. Surprisingly, we found that the gut microbiota may have a stronger association with *P. vivax* infection than with STH infection. One possible explanation is that STH co-infection could be affecting bacterial communities in a way that makes the host more permissive to *P. vivax* infection. Alternatively, *P. vivax* could be affecting the gut microbiota in a way that is prevented by STH infection. Longitudinal interventional studies that include deworming and anti-malarial treatment could provide greater causal evidence for these linkages.

## Methods

### Ethics statement

Written informed consent was obtained from parents or guardians of all study participants. Ethical approval was granted by the Committee on Human Ethics of the Health Sciences Department of the University of Córdoba, Monteria, Colombia.

### Study Design and Sample collection

Sample collection was performed between February and June 2019 municipality of Tierralta, Colombia. Tierralta is situated in the south-west of the department of Córdoba (8°10’ N, −76°04’E).

Children with *P. vivax* malaria were recruited at Hospital San José of Tierralta. *P. vivax* infection was diagnoses by a thick blood smear, performed in the morning. Children returned in the afternoon for diagnosis and, if positive, for treatment with Chloroquine (10 mg/kg, followed by 7.5 mg/kg at 24 and 48 h) and Primaquine (0.25 mg/kg for 14 days), according to the National Health Institute of Colombia guidelines. None of the children were hospitalized. Blood and stool samples were collected on the same day or the next day after *P. vivax* malaria diagnosis.

The groups of control children and STH-infected children were recruited at a local school, Institución Educativa Santa Fe de Ralito, within the municipality of Tierralta. The catchment area of the school is within the larger catchment area of the hospital. An informational meeting with the parents of children recruited for the study was held to explain the procedures and distribute screw-capped containers.Study participents returned the stool samples on the next day; at that time, study coordinators also collected blood samples.

### Collection and processing of stool samples

Samples were transported on the same morning to Universidad de Córdoba where they were divided into two portions: one portion was processed on the same day by Kato Katz and the other portion was stored at −20°C until DNA was extracted.

The species diagnosis and intensity of infection were determined by Kato Katz and results were recorded as egg per gram, following WHO guidelines (https://apps.who.int/iris/handle/10665/63821). Eggs per gram were recorded for *Trichuris trichiura, Ascaris lumbricoides* and hookworm. DNA was extracted using the DNeasy PowerSoil Kit (Qiagen) according to the manufacturer’s instructions.

### Microbiome sequencing and analysis

The 16S rRNA gene was amplified at the V4 hypervariable region and sequenced according to a multiplexing protocol described elsewhere (Caporaso et al. 2011; Caporaso et al. 2012) on the Illumina MiSeq system paired-end sequencing for 2 x 150bp reads. Automated 16s rRNA amplicon PCR libraries using GTC F/R primers were added to each sample; samples were then normalized and pooled. Microbiome bioinformatics were performed with QIIME 2 2019.4 (Bolyen et al. 2019). Raw sequence data were de-multiplexed and quality filtered using the q2-demux plugin and denoising with DADA 2 (Callahan et al. 2016). Amplicon sequence variants were aligned with mafft (Katoh et al. 2002) and used to construct a phylogeny with fasttree (Price et al. 2010). Alpha and beta diversity metrics were estimated using q2-diversity after samples were rarefied (subsampled without replacement) to 25,000 sequences per samples. Taxonomy was assigned to ASVs using the q2-feature-classifier (Bokulich et al. 2018) classify-sklearn naïve Bayes taxonomy classifier against the Greengenes 13_8 99% OTUs reference sequences (McDonald et al. 2012).

QIIME2 output files were imported into R using Phyloseq (McMurdie and Holmes 2013), and Jaccard distances between samples were calculated using non-metric multidimensional scaling (NMDS). A Principal Coordinate Analysis was done based on this dissimilarity matrix. Ellipses were superimposed, representing a 95% confidence level.

To examine the relative abundance of microbial taxa between comparison groups, we performed linear discriminant analysis (LDA) effect size (LEfSe) tests (Segata et al. 2011) with a p-value cutoff of 0.01, grouping samples by genus.

### Collection and processing of blood samples

Peripheral blood (5 ml in EDTA) was collected from each subject and processed on the same day by centrifugation at 1,372g for 5 min. Plasma was aliquoted and frozen at −80°C. Aliquots from all samples were shipped in dry ice to New York University School of Medicine. Cytokines were determined using Legendplex Inflammation panel I from Biolegend (San Diego, CA) for IL1-b, IL2, IL4, IL6, IL10, IL8, IL12p70, IL17A, IP10, TNFα, MCP1, IFNγ and TGF-β 1, on a FACSCalibur (Becton Dickinson, Franklin Lakes, NJ).

Peripheral blood was also collected in PAXgene Blood RNA tubes (PreAnalytiX, Qiagen, Valencia, CA) and stored at −80°C until RNA isolation. RNA isolation was performed using the PAXgene Blood RNA Kit (PreAnalytiX, Qiagen, Valencia, CA) according to the manufacturer’s instructions. RNA-Seq library preparation was done at the NYU School of Medicine Genome Technology Core using an automated stranded polyA enrichment library preparation protocol. Libraries were sequenced on a HiSeq 4000 (Illumina) with 2 x 50 cycles and for an average of 50 million reads per sample.

Raw RNA-Seq reads were aligned to the reference human genome Grch37 and the Ensembl reference transcriptome Homo_sapiens.GRCh37.82.gtf with Tophat (version 2.1.0), using all default parameter settings (Kim et al. 2013). Reads with a mapping quality score (MAPQ) of less than 30 and reads from mitochondrial DNA were removed. The number of filtered reads was subsequently counted for each gene by htseq-count, with the parameters—mode = union and—stranded = reverse (Anders et al. 2015). The resulting count matrix was used for downstream analyses.

### RNA-Seq data analyses

To perform PCA on the RNA-Seq data, we further filtered the above count matrix (where genes differential by sequencing batch were removed) and kept only genes with more than 10 counts in at least 10% samples. We next performed a variance stabilizing transformation (vst) on the filtered count data matrix generated as described above and removed genes with low variance (bottom half) using the varFilter function in the R/Bioconductor genefilter package (Gentleman et al. 2019). These filtering steps resulted in 6076 genes, on which we performed PCA using the vst-transformed count values.

We used DESeq2 (Love et al. 2014) to analyze differences between gene expression in comparison groups. Significance limits for upregulated genes were set as p<0.05, and the log2 fold change cutoff was set as greater than or equal to one. The genes identified by DESeq2 were then supplied to PANTHER; biological processes overrepresented in these genes relative to all genes in the *Homo sapiens* GO Ontology database (10/8/19) were enumerated, using Fisher’s Exact test for significance in PANTHER (Mi et al. 2019). The top ten specific subclasses (from hierarchically sorted output) were selected, based on lowest FDR.

### Random forest analyses

Metadata, clinical and cytokine test results were combined with 16S rRNA gene sequencing read counts (grouped by genus) and RNA-Seq read counts (filtering out all genes with less than 400 reads across all samples). Response variables used in separate analyses were: group (comparing one infection category to another), *P. vivax* parasitemia (continuous outcome) and *T. trichiura* egg count (continuous outcome). Variables were removed from the analyses that provided redundant information (for example, *T. trichiura* egg count was removed as a predictor when the response variable was *P. vivax* infected group versus the group co-infected with STH, since the observation of eggs in the stool was a determining characteristic for inclusion in the latter group). Analyses were done using randomForest in R (Liaw and Wiener 2002). For the group analysis, a classification tree was used with 10,000 trees, and 63 variables were tried at each split. When the response variable was *P. vivax* parasitemia, the random forest regression model based on 10,000 trees used 1346 variables at each split. The random forest regression model to predict *T. trichiura* egg count was based on 10,000 trees, and used 1345 variables at each split. Individual scatterplots were made based on variables of interest identified by these analyses. In order to add a best-fit line and r2 to the plot, a linear model was run.

### Data availability

Raw data of microbial 16S rRNA sequencing has been deposited in the European Nucleotide Archive with accession [insert here when available, currently processing]. Raw RNA-Seq data has been deposited on NCBI’s Gene Expression Omnibus with accession [insert here when available, currently processing].

---

**Figure S1:**
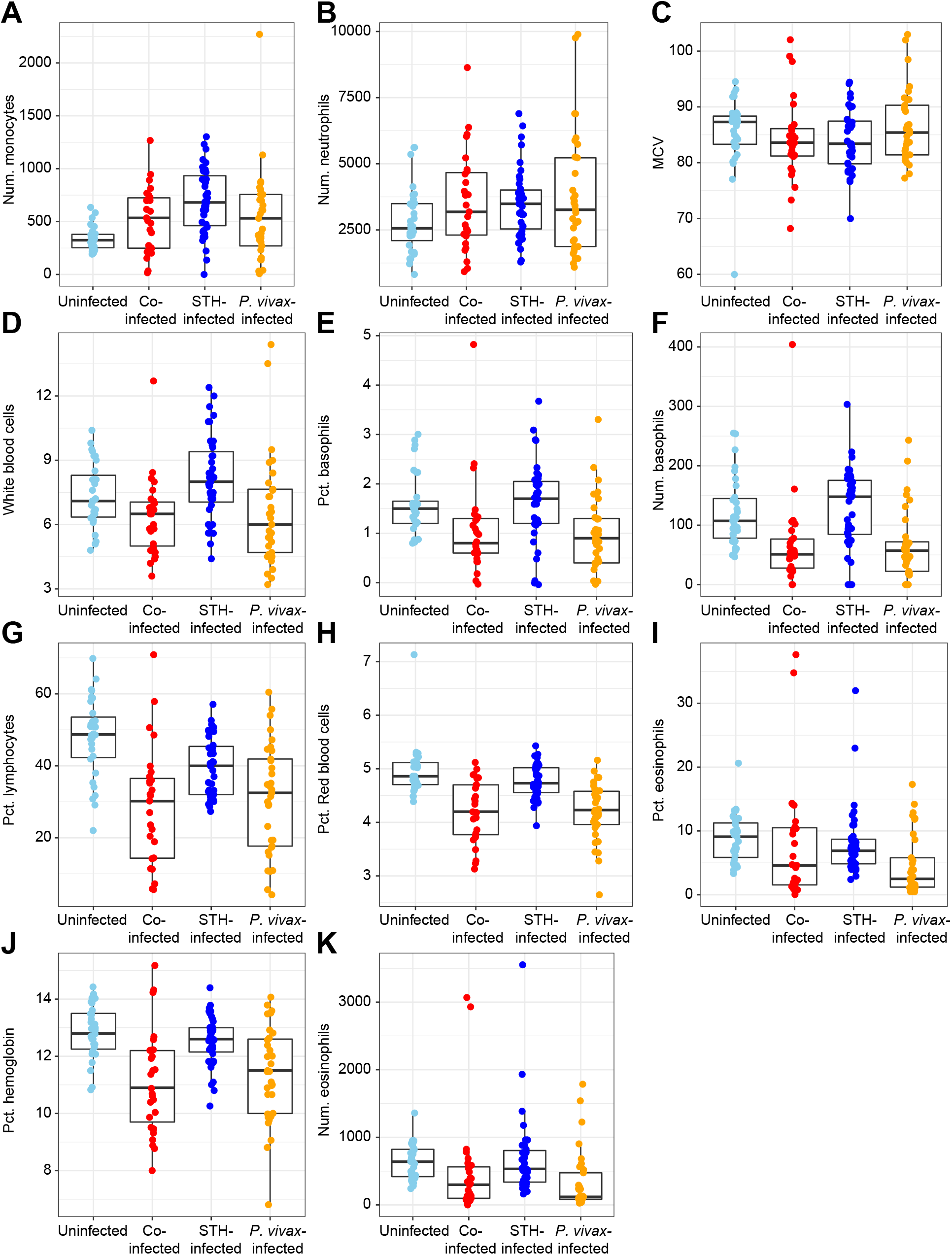
Additional boxplots of clinical data.

